# A new paradigm considering multicellular adhesion, repulsion and attraction represents diverse cellular tile patterns

**DOI:** 10.1101/2024.02.13.580045

**Authors:** Jose A. Carrillo, Hideki Murakawa, Makoto Sato, Miaoxing Wang

## Abstract

Cell sorting by differential adhesion is one of the basic mechanisms explaining spatial organization of neurons in early stage brain development of fruit flies. The columnar arrangements of neurons determine the large scale patterns in the fly visual center. Experimental studies indicate that hexagonal configurations regularly appear while tetragonal configurations can be induced in mutants. Mathematical models based on macroscopic approximations of agent based models (ARA models) are shown to produce a similar behavior changing from hexagonal to tetragonal steady configurations when medium range repulsion and longer-range attraction between individuals is incorporated in previous successful models for cell sorting based on adhesion and volume constraints. We analyse the angular configurations of these patterns based on angle summary statistics and compare between experimental data and parameter fitted ARA models showing that intermediate patterns between hexagonal and tetragonal configuration are common in experimental data as well as in our ARA mathematical model. Our studies indicate an overall qualitative agreement of ARA models in tile patterning and pave the way for their quantitative studies.

**2010 MSC:** 92C17, 92C37, 35Q92

## 1. Introduction

Organisms exhibit a variety of tile patterns, such as those seen in insect compound eyes, columnar structures in the brain, auditory epithelial cells, and liver lobules. Biological tile patterns often exhibit hexagonal patterns, which is often thought to be based on physical restrictions such as short circumference and high space-filling. However, aquatic arthropods such as shrimp exhibit tetragonal patterns in the compound eye [13]. It is also known that the compound eye of fruit fly, *Drosophila melanogaster*, normally exhibits a hexagonal pattern, but in some mutant backgrounds it changes to a tetragonal pattern [17]. Thus, organisms can produce either hexagonal or tetragonal tile patterns.

Columnar structures in the brain, which are formed by the cylindrical accumulation of multiple neurons, are the functional units of the brain, and their arrangement patterns are thought to play an important role in brain function. It is known that columnar structures in the fly visual center and microcolumns in the mouse cerebral cortex show a hexagonal arrangement, too [27, 28, 43]. However, the mechanisms controlling the tiling patterns of these columnar structures are unknown. Although hexagonal tile patterns tend to be preferred in the presence of physical constraints, it is also possible that the arrangement of the columns is not simply based on physical stability. If so, it is conceivable that the column arrangement may exhibit not only hexagonal but also tetragonal patterns.

Compared to the columns in the mammalian brain, which are composed of numerous neurons, fruit fly columns are structurally simpler, consisting of approximately 100 neurons. In addition, during early development (larval to early pupal stages), three neurons called R7, R8, and Mi1 play a central role in establishing the basic structure of the column: axons of R7 project to the center of the column and form a dot-like area; axons of R8 surround R7 and form a horseshoe-like circular area; axons of Mi1 surround R8 and occupy a grid-like region.

We found in [43] that the expression level of N-cadherin (Ncad), an evolutionarily conserved cell adhesion molecule, was strong in R7, intermediate in R8, and weakest in Mi1. The cells with stronger adhesion would be located on the inner side and the cells with weaker adhesion would be located on the outer side of the column. In fact, we have shown through a combination of experiments and a mathematical model that the cells are positioned according to the differences in cell adhesion, with R7 on the inner side, R8 just outside of it, and Mi1 at the outermost side [43, 9].

Cell sorting mechanisms have been considered in mathematical biology since the formulation of the differential adhesion hypothesis (DAH) by Malcolm Steinberg [35, 36, 37, 38] more than 50 years ago. Differential adhesion between different cell populations [20, 42] is now understood as a fundamental mechanism for cellular patterns, confirmed by experiments [10, 14, 15, 22, 44, 18, 19] and by mathematical models that are able to identify suitable parameters [21, 9, 34, 11, 12]. Mathematical Population Models (MPM) of cell sorting by differential adhesion are derived from Agent Based Models (ABM) by a coarse graining procedure usually referred as the mean-field approximation [8, 6, 9, 26]. MPMs include a nonlocal term incorporating long range attraction between cells. Long filopodia or protrusions are observed in experiments that produce contact forces between neighboring cells, see the discussion and derivation from Agent Based Models (ABM) in related contexts of zebra fish pattern formation and tissue morphogenesis [45, 7, 26]. These nonlocal population models can lead to aggregation-diffusion population models if volume effects are taken into account by introducing a strong repulsion at the origin. This approach leads to nonlinear diffusion terms [32, 9] in the population model instead of linear ones early used for this purpose [1]. Nonlinear diffusive terms allow for sharp interfaces between cell populations, being more natural than linear diffusion if population pressure is more important than Brownian motion fluctuations of the cells. In summary, previous works [1, 32, 9] introduced volume constraints by nonlinear diffusion and long range attraction by nonlocal terms with possible proliferation modelled by Fisher-KPP terms. Numerical experiments based on this MPM reported in [43] show that hexagonal type patterns for the columnar neuronal distribution in the fly visual center are ubiquitous for a large class of parameters.

However, our careful numerical exploration of MPMs for three populations densities (R7, R8 and Mi1) incorporating only differential adhesion via a long-range attractive potential does not lead to a transition between hexagonal to tetragonal tiling patterns. Let us mention that tetragonal configurations are proven to be the minimizers of the adhesion energy for repulsive-attractive interaction potentials [41], and numerically confirmed in [24]. From a biological perspective, it is known that ligands can lead to attractive or repulsive interaction at medium-range distances depending on the biochemical pathways involved [40, 39, 47].

In this study, we extend the previous differential adhesion model by incorporating a medium-range repulsion in between short-range and long-range attraction (ARA model) for the interactions between cells of the most cohesive population, the one with the strongest long-range attraction force, – R7 in the case of fly visual center – see Fig. 1. Symmetric differential adhesion for the other interand intracellular interactions is kept – R8 and Mi1 in the case of fly visual center. Using the ARA model, we have succeeded in reproducing hexagonal, tetragonal, and intermediate patterns interpolating between the two, matching the tailored-designed experimental results. Transitions in the tiling mechanism from tetragonal to hexagonal patterns is explained by varying the strength of the repulsive medium-range force and the length of the terrace between medium-range repulsion and long range attraction for the R7 population confirming the intuition derived from the theoretical results in [41]. We analyze this behavior using angle summary statistics to discern among hexagonal and tetragonal like patterns. Similar strategies based on summary statistics have been used in pattern classification for different purposes in mathematical biology [30, 31, 3, 2].

**Figure 1:**
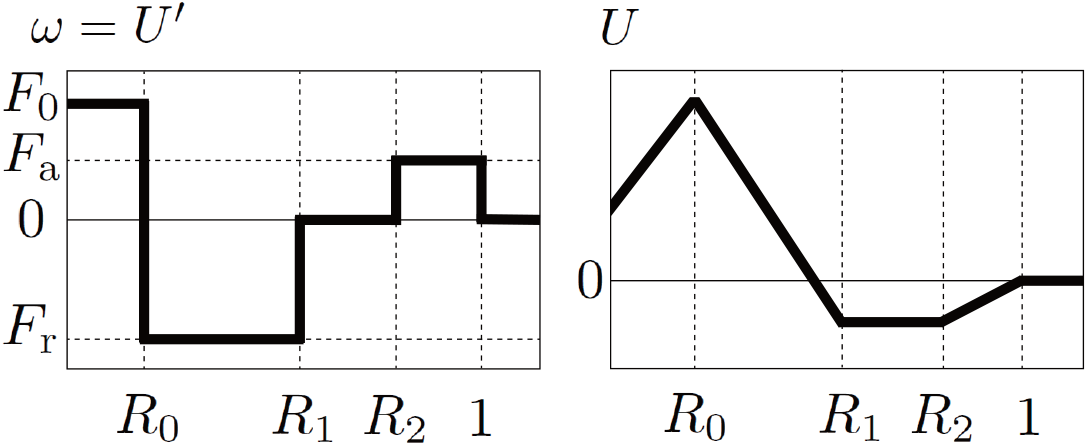
The shapes of the interaction kernel *ω*(*r*) and the corresponding potential *U* (*r*).

## 2. Materials and Methods

### 2.1 Cell-Cell adhesion basic model & Symmetry Patterns for a Single Population

As in [9], we introduce the population dynamics model for a single species in two spatial dimension [9] that reads as:

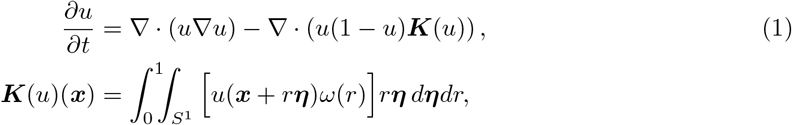

where *u*(***x***, *t*) denotes the population density at spacial position ***x****∈* R2 and time *t, S*^1^ is the unit circle and *ω* is a function that controls adhesion and repulsion between cells according to the distance from ***x***. The first term on the right-hand side of (1) denotes the basic behavior of cells, in which cells move from areas with high cell densities to areas with low cell densities due to population pressure. The integral term ***K***(*u*) of the second term represents that each cell perceives the situation within its own sensing radius *R* and moves in the desired direction accordingly. The sensing radius *R* is normalized to 1. The preference according to distance from itself is expressed by the interaction kernel *ω*. The term 1*− u* represents the effect of density saturation. We showed that the nonlinear diffusion, first term in (1), is crucial to have sharp interfaces instead of diffuse interfaces [1]. Moreover, this model is able to capture a good deal of mixing/sharp boundary behavior in tissue growth experiments [32]. Furthermore, Carrillo et al. [9] observed pattern formation by short-range adhesion and middle-range repulsion using (1). In particular, hexagonal spot patterns of cell populations have been ubiquitously obtained.

Theil [41] treated rigorously the geometric properties of the ground-state configuration of many-particle systems in two spatial dimensions under suitable assumptions on the interaction potential. He pointed out that the triangular lattice does not always have the lowest energy even for natural interaction potentials, and provided an interaction potential such that the energy per particle of the triangular lattice is higher than that of the square lattice. Based on his considerations, we use a simple potential. There are many reasonable choices for potentials to investigate biological phenomena of interest. In order to simplify the parameter estimation, in this paper, we deal with a piecewise linear potential *U* and hence the following piecewise constant kernel *ω* = *U* ^*′*^ as in Fig. 1 defined by:

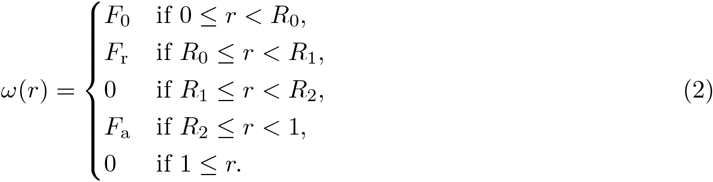

This kernel incorporates short-range adhesion (0 *≤ r < R*_0_), medium-range repulsion (*R*_0_ *≤ r < R*_1_), and long-range attraction (*R*_2_ *≤ r <* 1). The parameters *F*_0_ *>* 0, *F*_r_ *<* 0 and *F*_a_ *>* 0 control the strengths of cell-cell adhesion, that of repulsive and that of attraction, respectively. Motivated by Theil’s considerations, the terrace *U* (*r*) = const., i.e., *ω*(*r*) = 0 (*R*_1_ *≤ r < R*_2_) is introduced. Due to the balance of adhesion, repulsion, attraction and terrace, cells form small clusters, and the clusters are expected to arrange in a triangular or square lattice.

We illustrate numerically this behavior with different parameters for (1)–(2) when cells are initially gathered in one location. Throughout this work, all numerical simulations are carried out in a 2D domain [ *−* 2.8, 2.8)^2^ with the periodic boundary condition. The problems are discretized by the standard explicit upwind finite volume method [5], and the nonlocal terms are calculated by numerical integrations explained by Murakawa and Togashi [32]. The initial value is *u*_0_ = 0.1 inside a circle with a center at the origin and a radius of 0.2, and *u*_0_ = 0 otherwise. Grids where *u ≥*0.01 are drawn in green and grids where *u <* 0.01 are painted in white. Therefore, the green areas represent clusters of cells. Fig. 2 shows the numerical results with two sets of parameters.

**Figure 2:**
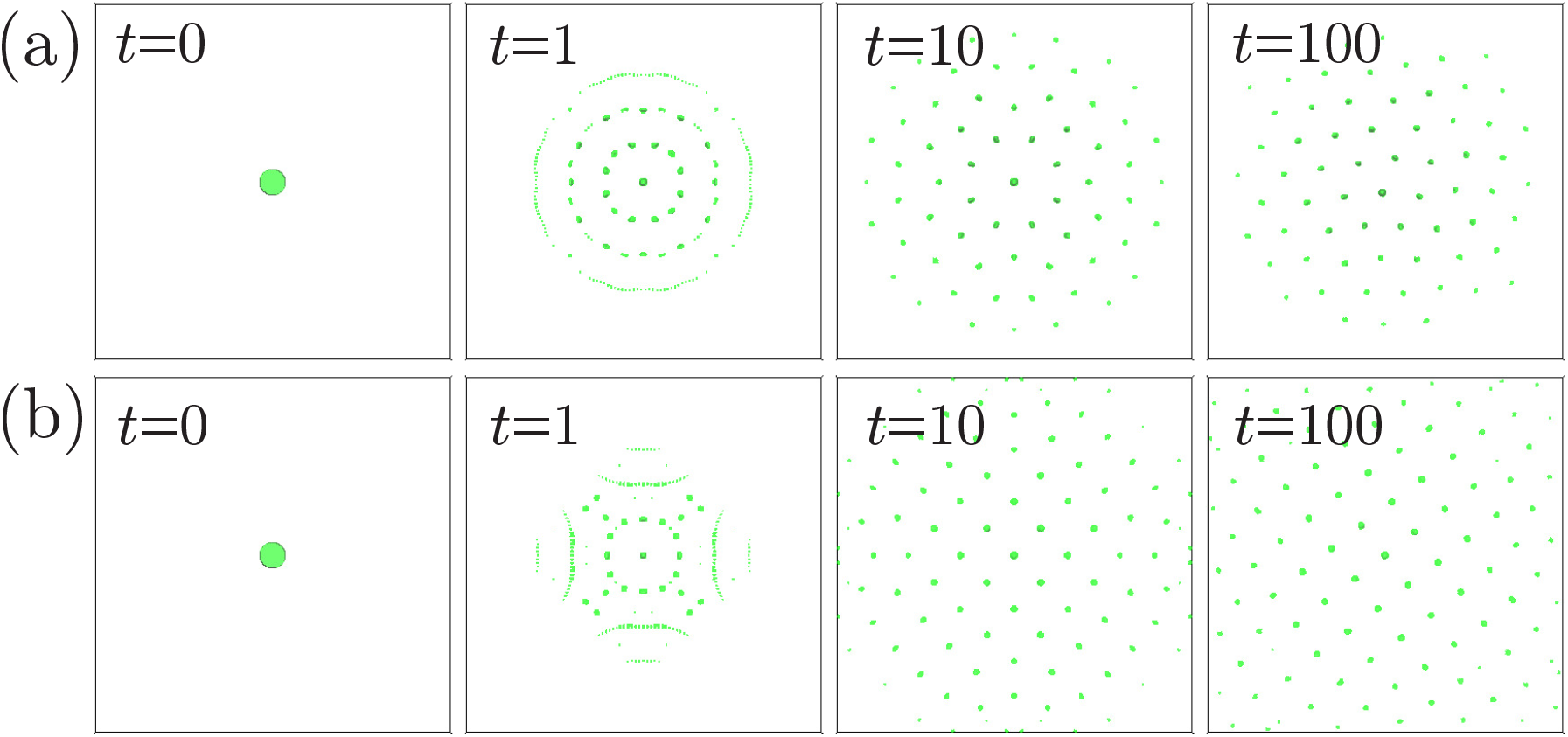
Numerical results for (1)–(2) with the following parameters. A: *R*_0_ = 0.14, *R*_1_ = 0.55, *R*_2_ = 0.58, *F*_0_ = 5000, *F*_r_ = *−*5000, *F*_a_ = 500. B: *R*_0_ = 0.14, *R*_1_ = 0.55, *R*_2_ = 0.88, *F*_0_ = 5000, *F*_r_ = *−*4000, *F*_a_ = 500.

We observe that cells gathered at the origin are scattered by medium-range repulsion, but form small clusters by short-range adhesion. The small clusters are arranged regularly due to the balance between medium-range repulsion and long-range attraction. In Fig. 2(a), cell clusters are arranged in a triangular lattice, and in Fig. 2(b), cell clusters are arranged in a square lattice. The main difference between the two is the size of the terrace, *i*.*e. R*_2_.

### 2.2 Experimental Data

The core columnar neurons, R7, R8, and Mi1, form concentric domains that establish the basic columnar structure in the larval brain, which grow into a mature three-dimensional column structureduring the pupal stage [43]. In the early larval stage, axons of R8, R7, and Mi1 sequentially project to the column, and in the late larval stage, they form a concentric column structure. In this study, the simulation starts when all axons of three types of core columnar neurons project to the column in the late larval stage. In the larval stage, the number of columns is still small and the quantitative results of the column arrangement pattern are not stable. So, we focus on the column arrangement pattern in the brain during the early pupal stage (20 hours after puparium formation: APF). In addition to R7, R8, and Mi1, many other types of neurons project to the columns, but for simplicity, only three types of columnar neurons were considered in this study.

Always centrally located in the column, R7 specifically expresses the cell adhesion molecule Fasciclin II (Fas2)[25, 46]. At 20 hours APF, brains were stained with mouse anti-Fas2 and Cy3-conjugated anti-mouse antibodies, mounted so that the column surface was placed horizontally to the cover glass [16]. The tips of the R7 axons were imaged using a confocal laser microscopy. After selecting data in which most of the columns were located in the same plane, individual R7 terminals were selected using the *Wand* function of Image J. The coordinates of their centers of mass were quantified using the *Measure* function, and these were used as the coordinates of the columns.

In addition to wild-type flies, we used a fly strain that expresses a transcription factor Gal4 in R7-specific manner (*R7-Gal4*), *UAS-Ncad RNAi* strain that knocks down Ncad under Gal4 control, and *UAS-Ncad* strain that overexpresses Ncad. Combining these flies, we artificially reduced and enhanced adhesion by Ncad in R7-specific manner and obtained column coordinates as above.

The distribution of the columns as visualized by Fas2 staining appeared to be hexagonal in control brains, see Fig. 3(a), whereas the pattern showed some irregularity when Ncad was either knocked-down or over-expressed in R7, see Fig. 3(b) and Fig. 3(c). To quantify the spatial pattern distribution of the columns, we shall formulate a quantitative method to analyze the symmetries of image data as explained below in the next section.

**Figure 3:**
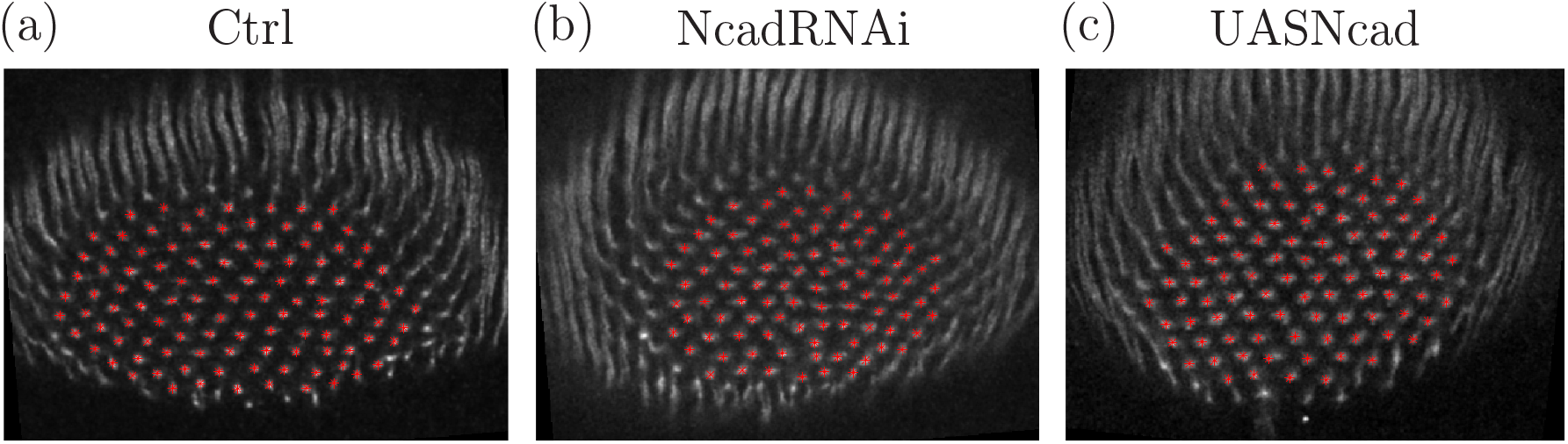
The columnar patters from the (a) control, (b) NcadRNAi and (c) UASNcad experiments. The columnar positions extracted from them are marked with red.

Although only R7 was visualized in biological experiments, individual columns also contain R8 and Mi1. Thus, we extended the single species model to the three-species model representing the densities of R7, R8, and Mi1 with *u, v*, and *w*, respectively, as described in [43, 9]. Our three-species mathematical model representing the densities *u, v, w* of R7, R8, and Mi1, respectively, is formulated as follows:

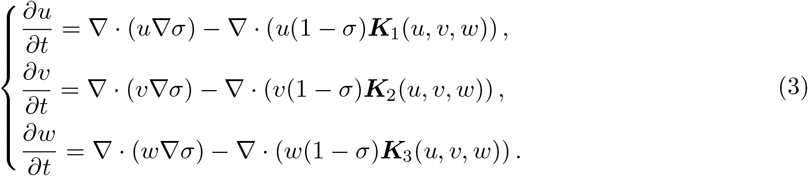

Here, the total density of neurons is denoted by *σ*= *u* + *v* + *w*. This Partial Differential Equations (PDEs) model includes volume size constraints in the form of total population pressure, nonlocal interactions incorporating attraction and repulsion, and density saturation effects. We refer to [9] for the full mathematical description. The nonlocal interactions ***K***_*i*_ (*i* = 1, 2, 3) will be fully specified in Section

### 2.3 Symmetry Indices & Data analysis

In order to analyze the symmetries on experimental and/or synthetic data, we introduce three symmetry indices. We first take a particle approximation both of the data and the simulations obtained with our continuum model. Centers of the neural columns in experimental data are directly identified in the corresponding images. The center of the local density peaks of R7 neurons in our simulations using the PDE model (3) is also directly obtained by its center of mass. Once we have this particle data approximation, we calculate the angle formed by each particle with its neighboring particles, and analyse the angle statistical distribution. There are various ways to determine angles of a home particle with neighboring particles, but we deal with the following three procedures:

- **NBH-1:** Consider the angle formed by the home particle and its 4 nearest neighbors. A slight perturbation of a perfect hexagonal pattern should produce angles close to those in Fig. 4(a), and the angle statistical distribution should concentrate at 60, 120, and 180 degrees as shown in Fig. 4(c). On the other hand, a slight perturbation of a perfect tetragonal pattern should only lead to angles like those in Fig. 4(b), and the angle statistical distribution should be concentrated at 90 degrees as shown in Fig. 4(d). The perturbations are generated by adding to 4096 particles in a perfect regular (hexagonal or tetragonal) configuration pattern a perturbation with different sizes. The 4 lines represent perturbations of 0.1%, 10%, 20%, and 30% of the maximal interparticle distance added to each particle.
- **NBH-2:** Consider the angle formed by the home particle and its 6 nearest neighbors.

**Figure 4:**
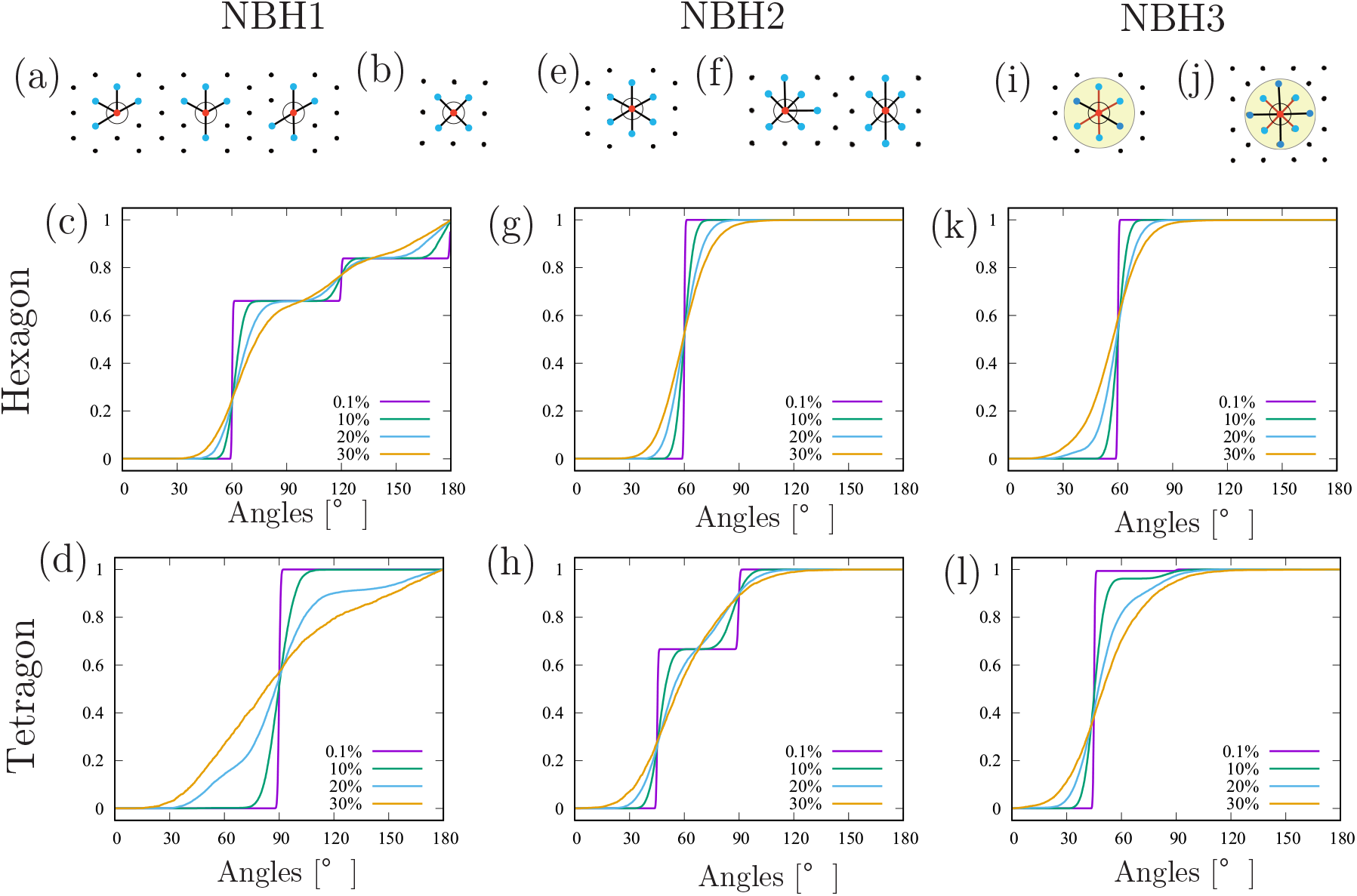
Sketch of three types of nearest neighborhoods, and cumulative angle distribution functions using these three types of nearest neighborhoods for perturbed hexagonal and tetragonal patterns. A slight perturbation of a perfect hexagonal pattern should only present angles close to those in Fig. 4(e). Therefore, the angles statistical distribution should be concentrated at 60 degrees as shown in Fig. 4(g). On the other hand, a slight perturbation of a perfect tetragonal pattern should only present angles close to those in Fig. 4(f), and the angle statistical distribution should concentrate around 45 and 90 degrees as shown in Fig. 4(h).
- **NBH-3:** Consider the angle formed by the home particle and the particles located in an circular region centered at the home particle. The circular region is specified by taking the average *d*^ave^ of the distances between the home particle and its 4 nearest neighbors. Consider all particles within a distance of 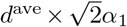 from the home particle. Then, calculate the angle formed between the home particle and each of these particles. In this paper, we set *α*_1_ = 1.1. Fig. 4(i) and (j) show examples for the hexagonal and tetragonal patterns, respectively. The lengths of the red bonds are the distance between the home particle and its four nearest neighbors, and the range is drawn in light green. For a slight perturbation of a perfect tetragonal pattern, the number of particles within the circular neighborhood should be 6, and the angle statistical distribution must concentrate at 60 degrees as shown in Fig. 4(k). On the other hand, for a slight perturbation of a perfect tetragonal pattern, the number of particles within the circular region should be 8, and the angle statistical distribution should be concentrated at 45 degrees as shown in Fig. 4(l).

The arrangement of the columns is analyzed by comparing the angle distribution function of experimental/numerical data with those of perturbations applied to regular lattices. The regular lattice discussed here is the one shown in Fig 5, and is a one-parameter family parameterized by *h*. It is the hexagonal lattice when 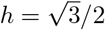, the square lattice when *h* = 1*/*2, and a centered rectangular lattice otherwise.

**Figure 5:**
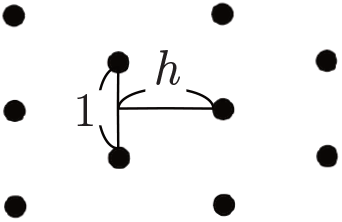
Scheme of the regular lattice and the parameter *h*.

Let 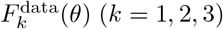 be the cumulative statistical distribution function of angles using NBH-*k* obtained from the position of the column based on experimental/numerical data, and let *F*_*k*_(*θ*; *h, p*) be the cumulative distribution function of angles of the perturbed regular lattice with a parameter *h* and a disturbance *p*%. We calculate the *L*^2^ error between 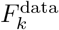 and *F*_*k*_(*·* ; *h, p*), and define the numerical pair (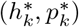) that minimizes this difference, namely,

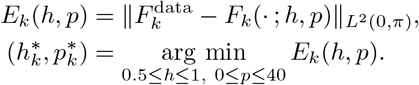

Let the value of 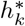 be denoted by Index-*k*.

Notice that since the number of particles is finite, a careful attention must be paid to the edges of the particle clusters. Calculate the distance *d*_*i*_ from each particle named *i* to its nearest neighboring particle and let the median of *d*_*i*_ be denoted as *d*. If there are 6 or more particles within a radius Of 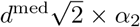 from a particle, that particle can be considered as a home particle. Otherwise, it is regarded as an edge particle. In this paper, we set *α*_2_ = 1.2.

## 3 Results

### 3.1 Analysis of symmetries on experimental data

We analyze five control (Ctrl), eleven Ncad knockdown (NcadRNAi) and eight Ncad over-expression (UASNcad) data sets using Indices-1,2,3. The results are summarized in Fig. 6. In order to demonstrate the reliability of the indices, we pick up data with index 3 close to the average value one by one from the Ctrl, NcadRNAi, and UASNcad datasets, and plot them in Fig. 3. We plot the cumulative distribution functions of the angles corresponding to NBH-1–3 in Fig 7. The two lines represent the cumulative distribution functions of the angles 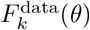obtained from experimental data and 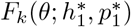of a perturbed regular lattice fitted to the experimental data. Index-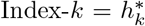, the corresponding 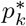 and error 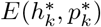 are summarized in Table 1. Everything fits well, and those indices appears to be useful.

**Table 1:** The parameters and errors 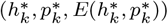 used in Fig. 7.

|  | Ctrl | NcadRNAi | UASNcad |
| --- | --- | --- | --- |
| Index-1 | (0.5735, 0.19, 0.006781) | (0.6157, 0.1872, 0.009073) | (0.64795, 0.1764, 0.005705) |
| Index-2 | (0.5962, 0.1864, 0.007195) | (0.6135, 0.204, 0.004349) | (0.6502, 0.1908, 0.004191) |
| Index-3 | (0.57645, 0.1788, 0.006575) | (0.6162, 0.2152, 0.005594) | (0.65065, 0.1764, 0.00632) |

**Figure 6:**
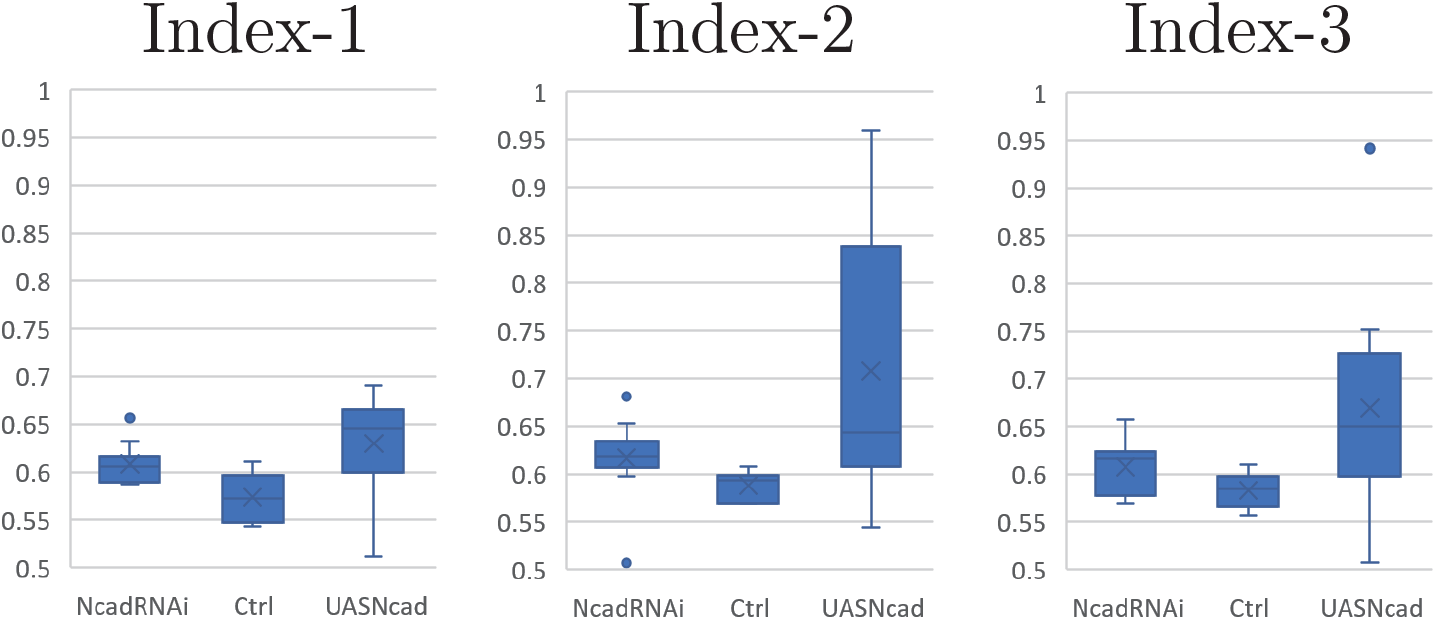
Boxplots of Indices-1,2,3 from 5 Ctrl, 11 NcadRNAi and 8 UASNcad experimental data sets.

**Figure 7:**
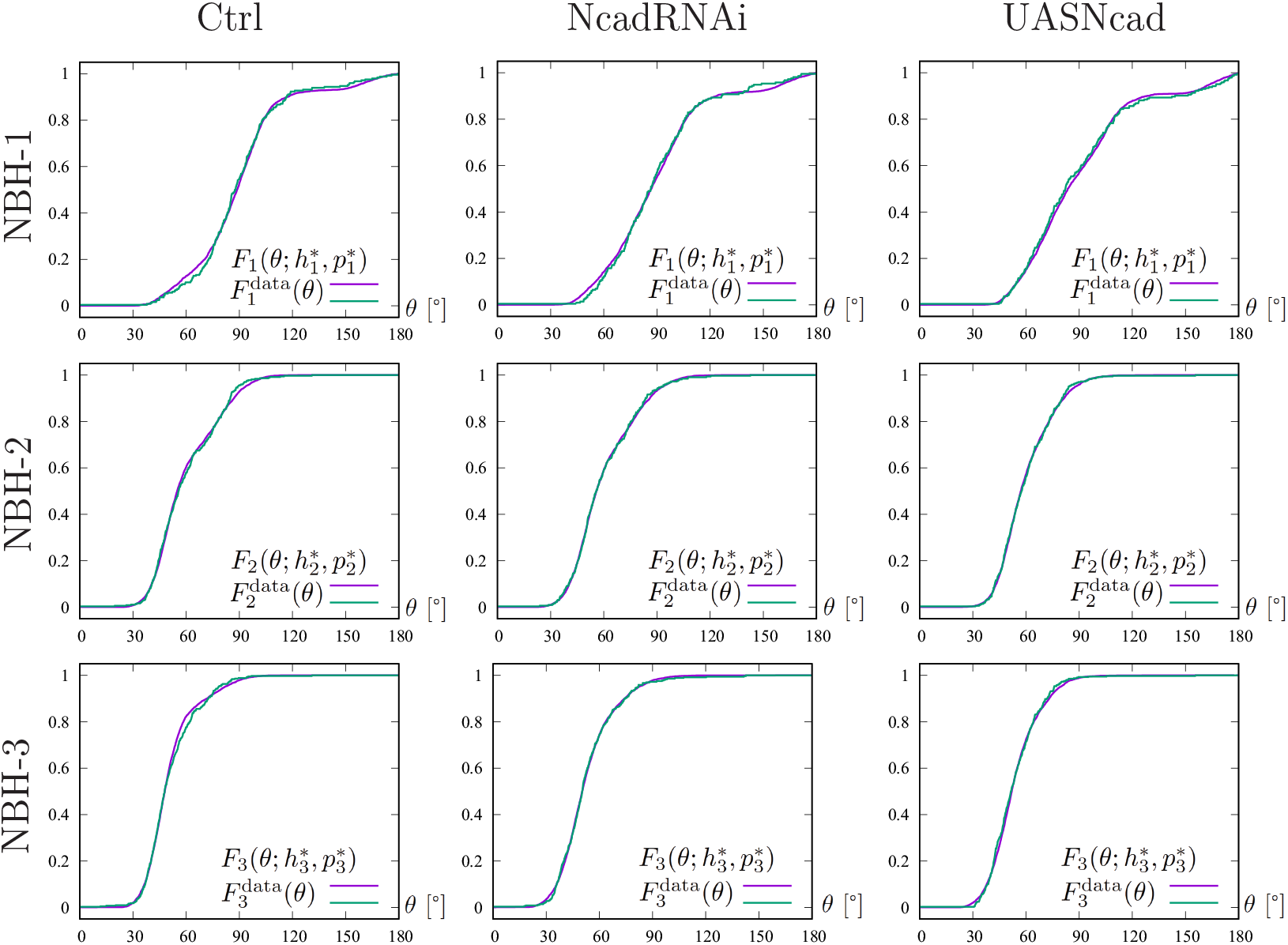
The cumulative distribution functions of the angles of the columnar patters from the control, NcadRNAi and UASNcad experiments in Fig. 3 corresponding to NBH-1–3, respectively. The two lines represent 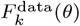 obtained from experimental data (green) and 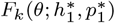 a perturbed regular lattice fitted to the experimental data (magenta).

In contrast to our initial naive impression that the distribution of the columns is hexagonal in control brains, see Fig. 3, the results of our symmetry analyses shown in Fig. 6 indicate that the pattern is always at an intermediate state between hexagonal and tetragonal. Furthermore, the control pattern is rather tetragonal (lower indices) while the patterns found in NcadRNAi and UASNcad conditions are rather hexagonal (higher indices). We next asked if a similar transition of distribution patterns can be reproduced in numerical simulations.

### 3.2 Numerical simulations

In order to investigate how the strength of the attraction/repulsion affects the column arrangement, we carry out numerical experiments based on the ARA mathematical model (3) where the nonlocal interactions ***K***_*i*_ (*i* = 1, 2, 3) are given by

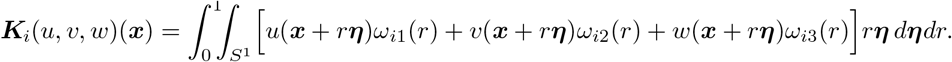

Since only the adhesion of R7 was artificially manipulated in the experiment, the ARA kernel is employed only for the interaction between R7 cells, while the cell adhesion kernels are used for other interactions, namely,

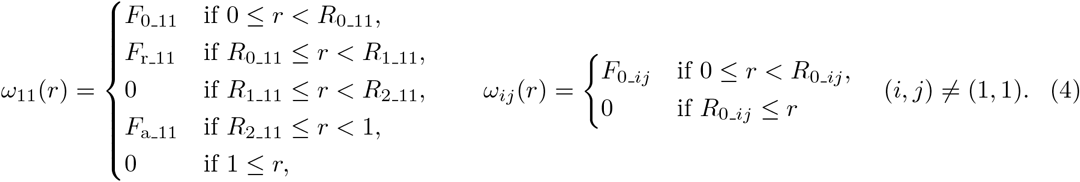

Numerical simulation using the larval column distribution pattern as the initial condition is difficult because the entire tissue grows during the larval stage and the number of columns itself increases. Therefore, we used the column coordinates of wild type flies at 20 hours APF as the initial conditions.

There are many parameters in the kernel (4), but our purpose is to investigate the effect of attraction on column arrangement. Our previous study revealed long cellular protrusions (filopodia) emanating from the columnar neurons such as R7 [41]. Ncad located at the tip of filopodia may mediate a long range attraction. Therefore, we vary *F*_a 11_ and *R*_2 11_ while keeping other parameters fixed as follows: *F*_0 11_ = 2000, *F*_r 11_ = *−*2000, *R*_0 11_ = 0.14, *R*_1 11_ = 0.55, *F*_0 22_ = 25, *F*_0 12_ = *F*_0 21_ = 50, *R*_0 12_ = *R*_0 21_ = 0.3, *R*_0 *j*2_ = *R*_0 2*j*_ = 0.14, *F*_0 33_ = 10, *F*_0 13_ = *F*_0 31_ = 20, *F*_0 23_ = *F*_0 32_ = 15, *R*_0_ *j*3_ = *R*_0 3*j*_ = 0.14. To determine that the numerical solution has approached the steady state, the computation was stopped if the *l*^2^(Ω) error between the numerical solutions at time *t* and at *t* 1 falls below a tolerance 10^*−*5^.

Numerical calculations are performed from the experimental column coordinate, that is shown in Fig. 8(a), for each parameters *F*_a 11_ and *R*_2 11_. The indices of the columnar coordinates are calculated when sufficient time has passed, and the results are depicted in Fig. 8(b). The values of the indices are displayed in colors. Additionally, if the columnar pattern is significantly disrupted, it is painted in white. The numerical columnar coordinates for several selected parameters are displayed in Fig. 8(c)– (g). Highly organized tetragonal pattern is observed when (*F*_a 11_, *R*_2 11_) is (850, 0.85), see Fig. 8(g). On the other hand, in the vicinity of *R*_2 11_ = 0.64, differences in the indices can be observed due to variations in *F*_a 11_. Even though the indexes are different, their appearance does not change significantly, see Fig. 8(c)–(e). It becomes clear that a quantitative perspective is necessary. When the attraction is too strong, it can be observed that the columnar structure breaks down, see Fig. 8(f). Fig. 8(h) shows the details of the change in the indices when *F*_a 11_ is changed while *R*_2 11_ = 0.64 is fixed. It is assumed that *F*_a 11_ between 296 and 305 corresponds to the control state, and boxplots are made in Fig. 8(i) including cases where the attractions are weak (*F*_a 11_ = 251–260) and strong (*F*_a 11_ = 411–420). In Fig. 8(h), calculations are performed by incrementally changing the value of *F*_a 11_ by one. From these data, we found a location where the change in Index-1 with respect to *F*_a 11_ was convex, and selected a range of width 10 as corresponding to each of NcadRNAi, Cntl, and UASNcad. Although those are based on artificial numerical experiments and the results depend on the initial states, it is observed that the pattern deviates from the tetragonal pattern due to suppression and over-expression of attraction forces similar to the results in Fig. 6. It is technically difficult to measure real parameters such as *F*_a 11_, and there is no guarantee that the parameters we found above directly correspond to the real phenomenon. Nevertheless, the method we developed opens an avenue to study biological tiling patterns in a quantitative manner.

**Figure 8:**
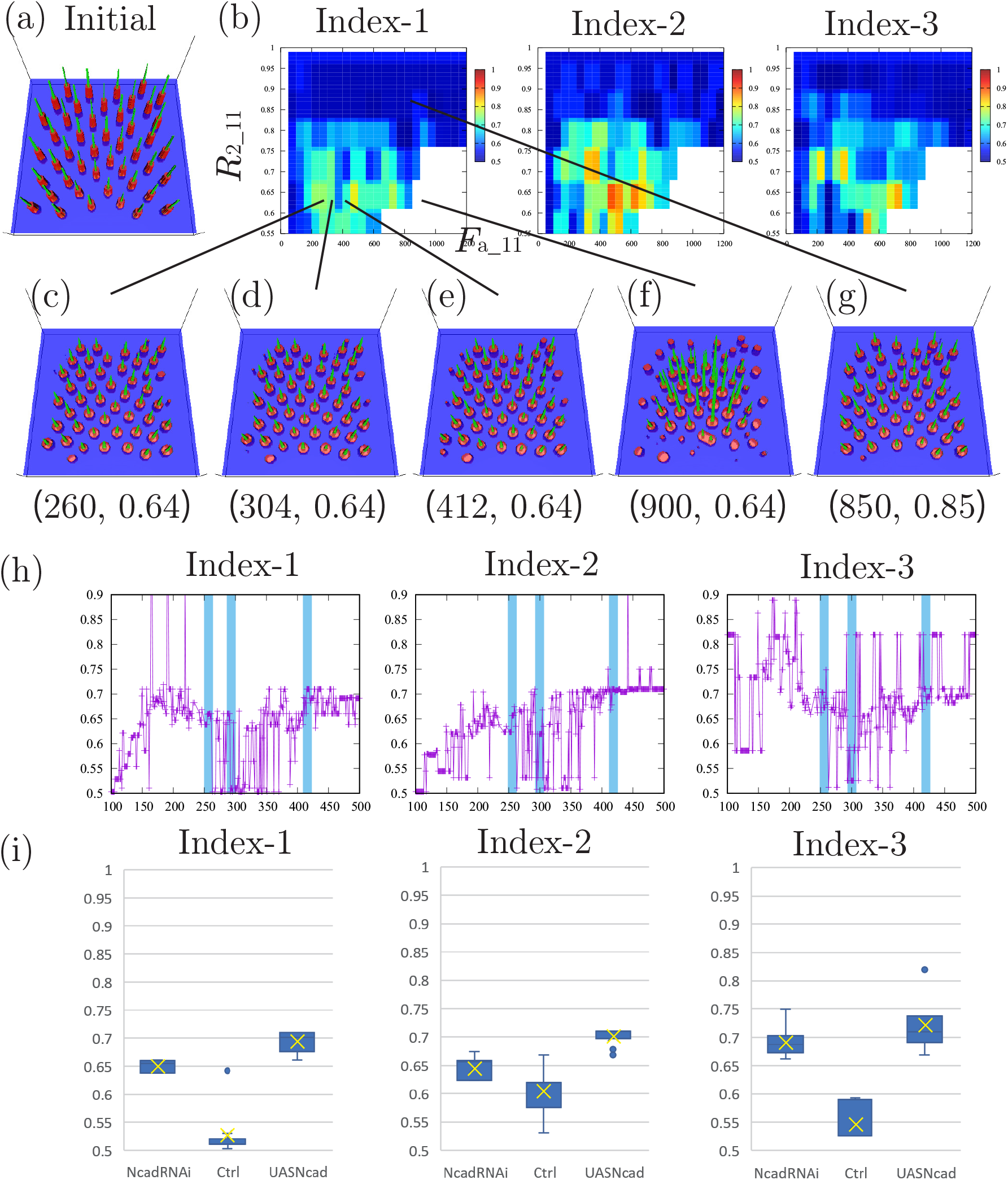
(a) Initial state based on experimental data. (b) Indices using different parameters *F*_a 11_ and *R*_2 11_ at which numerical solutions were reached numerical steady states. (c)–(g) Several selected numerical steady states. (h) Indices using different parameters *F*_a 11_ with fixed *R*_2 11_ = 0.64 at which numerical solutions were reached numerical steady states. (i) Boxplots of Indices-1,2,3 from *F*_a 11_ = 251–260 (labeled NcadRNAi), *F*_a 11_ = 296–305 (labeled Ctrl) and *F*_a 11_ = 411–420 (labeled UASNcad) numerical data sets. These parameter ranges are shown in light blue in (h). Each average value is indicated by a cross, and in Figs. (c) to (e), results using data close to each average value are plotted.

## 4. Discussion

A wide variety of tile patterns are found in a variety of tissues and are not simply hexagonal or tetragonal. Intermediate patterns between hexagonal and tetragonal may also be common, as seen in the columns in the fly brain.

The arrangement of the columns may play a role in visual information processing in the fly visual system, which includes the compound eye and visual center in the brain. The fly visual system computes motion of visual objects by temporally comparing contrast signals between neighboring points in space received by eye units in the compound eye. The arrangement of eye units in the compound eye and the columnar structure in the brain must be linked to visual information processing. In the mouse cerebral cortex, arrangement of the microcolumns roughly show hexagonal arrangement [27]. However, their arrangement was not rigorously quantified. Careful quantification and functional study of their arrangement are required.

In this study, we developed the ARA model to deal with variation of the tile patterns found in the fly visual center. The ARA incorporates medium-range repulsion embedded between shortand long-range attraction, and reproduces different tile patterns composed of multiple species (or multiple cell types). We focused on the function of a cell adhesion molecule, Ncad, which is thought to mediate contact-mediated adhesion, or short-range attraction. However, our previous study revealed long cellular protrusions (filopodia) emanating from the columnar neurons such as R7 [43]. Considering that cell adhesion can be mediated by Ncad located at the tip of filopodia, Ncad-mediated adhesion may occur at a long range and is compatible with the long-range attraction, which is related to the parameter *F*_a 11_ in the numerical experiments performed in this study.

The other concern is the nature of medium-range repulsion. Slit, Semaphorin, Wnt, and Netrin are typical families of diffusible repulsive molecules [40, 39, 47]. Dscam is a family of cell adhesion molecules that induces contact-mediated repulsion, which may also act at a long range through the filopodia [23, 29, 33]. Netrin has been proposed to switch between attractant and repellent depending on the receptor configuration [39]. Our study indicates the need of identifying the molecular basis of medium-range repulsion that acts on the arrangement of the columns in the fly visual center.

In addition, we have developed a quantitative method to analyse the symmetric distribution patterns of cellular units such as columns. The same method may be used to quantify the distribution pattern of microcolumns in the mouse cerebral cortex [27]. Here, we emphasize that finding alternative indices for angle statistics to improve the data analysis of hexagonal versus tetragonal tile patterns is a timely research topic. Similar strategies for fitting parameters using summary statistics, pattern simplicity scores or persistent-homology approaches have been used to characterize pattern formation in several mathematical biology models, see [30, 31, 4, 3, 2, 26] for instance. Our study provides a mathematical basis for studying a wide range of tile patterns found in biological and non-biological systems.

## Author Contributions

**Conceptualization:** Jose A. Carrillo, Hideki Murakawa, Makoto Sato.

**Data curation:** Hideki Murakawa, Makoto Sato.

**Formal analysis:** Jose A. Carrillo, Hideki Murakawa, Makoto Sato.

**Funding acquisition:** Jose A. Carrillo, Hideki Murakawa, Makoto Sato, Miaoxing Wang.

**Investigation:** Jose A. Carrillo, Hideki Murakawa, Makoto Sato, Miaoxing Wang.

**Methodology:** Jose A. Carrillo, Hideki Murakawa, Makoto Sato.

**Project administration:** Jose A. Carrillo, Hideki Murakawa, Makoto Sato.

**Resources:** Hideki Murakawa, Makoto Sato.

**Software:** Hideki Murakawa.

**Supervision:** Jose A. Carrillo, Hideki Murakawa, Makoto Sato.

**Validation:** Hideki Murakawa, Makoto Sato, Miaoxing Wang.

**Visualization:** Hideki Murakawa, Makoto Sato.

**Writing - original draft:** Jose A. Carrillo, Hideki Murakawa, Makoto Sato.

**Writing - review & editing:** Jose A. Carrillo, Hideki Murakawa, Makoto Sato.

